# Antidiabetic effect of 1-deazapurines derivatives with alpha-glucosidase Enzyme: A molecular tools approach

**DOI:** 10.1101/2024.02.20.581286

**Authors:** Faiza Boukli Hacene, Sabri Ahmed Cherrak, Wassila Soufi, Said Ghalem

## Abstract

Alpha-glucosidase inhibition has been shown by several 1-deazapurine derivatives that have already been synthesized and evaluated in vitro, providing a potential treatment target for type 2 diabetes. Six of them were shown to have higher percentages of inhibition against the alpha-glucosidase enzyme and lower IC50 values. The binding mechanism and stability of the generated complexes are investigated in the current work using various molecular modeling methodologies. The ligands L17, L11, and L4 present the best binding energy with the establishment of interaction toward the site active, based on the results of the molecular docking simulation. The stability of the chosen complexes was then validated using molecular dynamics simulation. However, ADME-T prediction and Drug-likeness results show that these compounds have promising pharmacokinetic properties and oral bioavailability. Finally, these results imply that compounds L4 and L11 are very promising as a target for creating a lead molecule for type 2 diabetes.

## Introduction

Type 2 diabetes mellitus (T2DM) is the most prevalent form of diabetes, accounting for at least 90% of all cases **[1–3]**. It characterized by chronically elevated blood glucose levels (hyperglycemia) which will cause many complications including neuropathy, nephropathy along with retinopathy, and cardiovascular disorders **[4,5]**.

Alpha-glucosidase inhibitors (AGIs) are oral antihyperglycemic drugs often given to persons with T2DM to control postprandial blood glucose (PPG) levels **[6,7]**. They inhibit the upper gastrointestinal enzymes that convert complex carbohydrates into glucose **[8]**.

Inhibition of alpha-glucosidase in the gastrointestinal tract delays carbohydrate digestion, leading to a reduction in the rate of glucose absorption and, consequently, postprandial blood glucose and insulin level **[9]**, which is considered a viable prophylactic treatment for hyperglycemia. The overall efficacy of this class of drugs appears high: AGIs (Acarbose, Miglitol and Voglibose) can be used safely in the elderly by decreasing the risk of hypoglycemia and toxicity **[8]**. Acarbose has been shown to reduce the risk of cardiovascular diseases such as systolic blood pressure **[10]**; major atherosclerotic events and ischemic stroke **[11]**, but the biggest limiting factor for wider use of these drugs is the side effects such as diarrhea, abdominal distention, and flatulence **[12]**.

1-deazapurine derivatives represent a significant class of nucleic heterocyclic found in numerous molecules within the field of medicinal chemistry **[13,14]**. Previous studies has suggested that 1-deazapurines and derivatives exhibit several biological activities which showed excellent inhibitory potency in vitro against alpha-glucosidase **[15,16]**.

A series of 1-deazapurine derivatives structures have been selected for a study involving molecular docking and molecular dynamics using both (MOE software and GROMACS software) **[17–19]**. Our focus is on comparing these inhibitors with the anti-diabetic drug Acarbose. In this context, docking studies primarily aim to predict the affinity between the ligand and the active site residues of the protein receptor, aiding in determining a reasonable conformation **[20]**. Additionally, the study involves predicting the in silico ADMET properties of the compounds for drug likeness and bio-compatibility.

Furthermore, the investigation aims to validate the stability of the most potent inhibitors against alpha-glucosidase through molecular dynamics simulations.

## Material and methods

### Biological Data

A series of 1-deazapurines derivatives synthesized **[21–23]** have been tested in vitro for enzyme inhibition alpha glucosidase **[15]**, the reported experimental inhibition activities are listed in Table 1. The activities corresponding to the 19 ligands mentioned above were given for their IC50 and their percentage of inhibition.

**Table 1.**
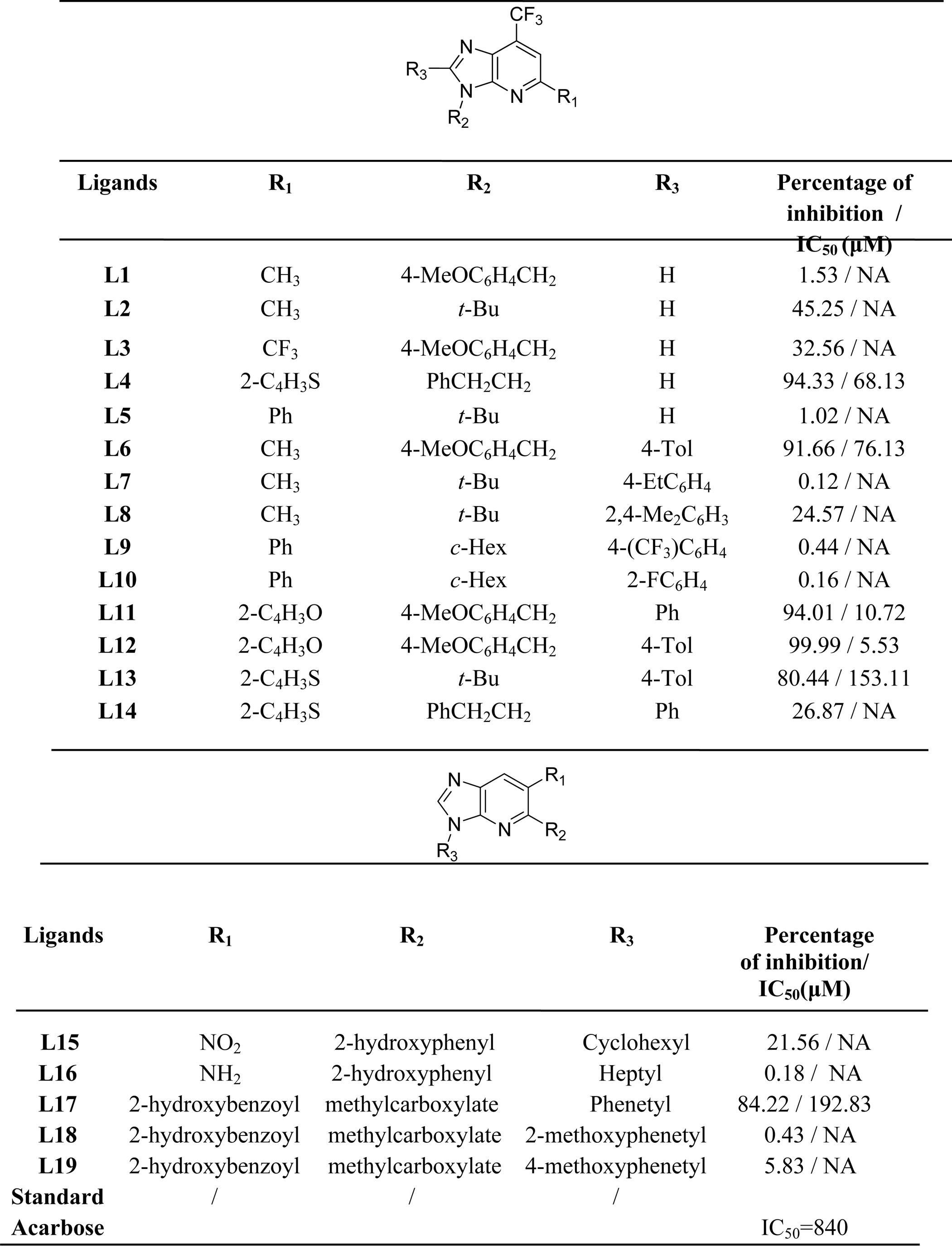
Structure of ligands and their respective percentage of inhibition and IC_50_ [15].

|  |  |  |  |  |
| --- | --- | --- | --- | --- |
| Ligands | R<sub>1</sub> | R<sub>2</sub> | R<sub>3</sub> | Percentage of inhibition /<br>IC<sub>50</sub> (μM) |
| L1 | CH<sub>3</sub> | 4-MeOC<sub>6</sub>H<sub>4</sub>CH<sub>2</sub> | H | 1.53 / NA |
| L2 | CH<sub>3</sub> | *t*-Bu | H | 45.25 / NA |
| L3 | CF<sub>3</sub> | 4-MeOC<sub>6</sub>H<sub>4</sub>CH<sub>2</sub> | H | 32.56 / NA |
| L4 | 2-C<sub>4</sub>H<sub>3</sub>S | PhCH<sub>2</sub>CH<sub>2</sub> | H | 94.33 / 68.13 |
| L5 | Ph | *t*-Bu | H | 1.02 / NA |
| L6 | CH<sub>3</sub> | 4-MeOC<sub>6</sub>H<sub>4</sub>CH<sub>2</sub> | 4-Tol | 91.66 / 76.13 |
| L7 | CH<sub>3</sub> | *t*-Bu | 4-EtC<sub>6</sub>H<sub>4</sub> | 0.12 / NA |
| L8 | CH<sub>3</sub> | *t*-Bu | 2,4-Me<sub>2</sub>C<sub>6</sub>H<sub>3</sub> | 24.57 / NA |
| L9 | Ph | *c*-Hex | 4-(CF<sub>3</sub>)C<sub>6</sub>H<sub>4</sub> | 0.44 / NA |
| L10 | Ph | *c*-Hex | 2-FC<sub>6</sub>H<sub>4</sub> | 0.16 / NA |
| L11 | 2-C<sub>4</sub>H<sub>3</sub>O | 4-MeOC<sub>6</sub>H<sub>4</sub>CH<sub>2</sub> | Ph | 94.01 / 10.72 |
| L12 | 2-C<sub>4</sub>H<sub>3</sub>O | 4-MeOC<sub>6</sub>H<sub>4</sub>CH<sub>2</sub> | 4-Tol | 99.99 / 5.53 |
| L13 | 2-C<sub>4</sub>H<sub>3</sub>S | *t*-Bu | 4-Tol | 80.44 / 153.11 |
| L14 | 2-C<sub>4</sub>H<sub>3</sub>S | PhCH<sub>2</sub>CH<sub>2</sub> | Ph | 26.87 / NA |
| Ligands | R<sub>1</sub> | R<sub>2</sub> | R<sub>3</sub> | Percentage of inhibition/<br>IC<sub>50</sub>(μM) |
| L15 | NO<sub>2</sub> | 2-hydroxyphenyl | Cyclohexyl | 21.56 / NA |
| L16 | NH<sub>2</sub> | 2-hydroxyphenyl | Heptyl | 0.18 / NA |
| L17 | 2-hydroxybenzoyl | methylcarboxylate | Phenetyl | 84.22 / 192.83 |
| L18 | 2-hydroxybenzoyl | methylcarboxylate | 2-methoxyphenetyl | 0.43 / NA |
| L19 | 2-hydroxybenzoyl | methylcarboxylate | 4-methoxyphenetyl | 5.83 / NA |
| Standard<br>Acarbose | / | / | / | IC<sub>50</sub>=840 |

### Enzyme and ligands preparation

The alpha-glucosidase structure (code 3L4Y) was retrieved from the Protein Data Bank (https://www.rcsb.org/) via X-ray diffraction, providing a three-dimensional structure at a resolution of 1.80 Å and the missing amino acids were optimized using modeller in UCSF chimera **[24]**. The inhibitors utilized in this study are a series of 1-deazapurine derivatives, as detailed in Table 1 **[21–23]**. These inhibitors have already been synthesized.

The construction and optimization of the ligands were conducted using the Molecular Operating Environment (MOE) software **[17]**.

### Molecular Docking

The Molecular Operating Environment (MOE) **[17]** was used to perform molecular docking calculations. Prior to this, all crystallographic structures (X-ray) of the chosen targets were made simpler by eliminating water molecules, ions, cofactors, and co-crystal ligands from their PDB structures. Using MOE software the active site of the enzyme was selected and energy minimization was realized for both enzyme and the inhibitors. Energy minimizing was done under the following conditions: temperature = 300°K, pH = 7, the geometry was performed using the field strengths in the MMFF94x implanted in MOE and Hamiltonian AM1.

### Pharmacokinetics/drug-likeness prediction

The number of drugs that failed in clinical trials due to poor absorption, distribution, metabolism, excretion, toxicity (ADMET) qualities has significantly decreased as a result of early stage drug development taking ADMET features into account **[25]**. Therfore pkCSM server (http://biosig.unimelb.edu.au/pkcsm/prediction) **[26]** was used for predicting and optimizing small-molecule pharmacokinetic and toxicity properties of the following parameters: blood–brain barrier (BBB), gastrointestinal absorption (GI), P-glycoprotein (P-gp) substrate, penetration, and cytochrome enzyme (CYP) inhibition. In addition the drug-likeness prediction is based on several criteria such as Lipinski’s rule of five **[27]**, Ghose’s rule **[28]**, Veber’s rule **[29]**, Egan’s rule **[30]**, Muegge’s rule **[31]** were performed from SwissADME (http://www.swissadme.ch/).

### Molecular dynamics (MD) simulation

We have access to a potent toolkit through molecular dynamics (MD) simulation, which allows us to follow and comprehend structure and dynamics in great detail literally, on scales where individual atoms can be tracked **[32]**. In this case study, we used MD simulations to perform stability tests against alpha-glucosidase utilizing the best-docked complex of 19 ligands.

CHARMM36 force field installed on a Linux system was used to execute molecular dynamics simulations using the GROMACS 2020 package **[18,19]**. The pdb2gmx program was utilized to prepare the protein topology. The protein complexes were solvated in a triclinic box. The steepest descent algorithm was used to minimize the system’s energy **[33]**. Four Na+ ions were added to the system to neutralize it after the protein was solvated using the TIP3 solvent model.

To stabilize the system, an overall pressure and temperature of 1 bar and 300 K were applied with a 2 fs time interval. The rescale, an enhanced Berendsen thermostat temperature coupling method, was utilized to maintain a consistent temperature inside the box. For NPT equilibration, the Parrinello-Rahman pressure coupling approach was applied. Using the steepest descent approach for 50,000 iteration steps and a cut-off up to 1000 kJ.mol-1 to lessen steric conflicts, the system’s energy was minimized. With a 1.0 nm Coulomb cut-off, long-range electrostatic interactions were treated using particle mesh Ewald (PME). A final 100 ns molecular dynamic run was performed on the system for every 2 fs step. The plot of the potential energy variations as a function of time (ps) was done with OriginePro **[34]**.

## Results and discussion

### Molecular docking

Molecular docking is used to predict the most likely 3D conformations of Small-molecule ligands within target binding sites, which also offers quantitative estimates of the energy fluctuations involved in the intermolecular recognition event **[35]**. The two separate processes of molecular docking are estimate of the binding energy for each expected conformation and exploration of the ligand conformational space within the binding cavity **[36,37]**.

### Score energy

The table **1** shows the presence of inhibitory activity for Acarbose and only for ligands L4, L6, L11, L12, L13 and L17, it is due to their substituted phenyl groups **[15,38]**.

Docking calculation results of the energy score are listed in the table 2. The complex17 (L17_3L4Y) shows the lowest binding energy with −6.1247 kcal/mol (highest affinity). The others complexes have a score energy ranging between −3.6741 and −5.9214. We note that the score energy of complexes co-crystallized_ligand_3L4Y and Acarbose_3L4Y have respectively −5.8036 and −5.6234 kcal/mol. Our molecular docking results are according to their inhibitory activity against alpha-glucosidase **[15]**.

**Table 2.** Energy score of 19 ligands of 1-deazapurines.

| Enzyme alpha-glucosidase (3L4Y) |  |  |  |  |  |  |  |
| --- | --- | --- | --- | --- | --- | --- | --- |
| Native Ligand rmsd 1.3288 |  |  |  |  |  |  |  |
|  | s | rmsd_refine | E_conf | E_place | E_score1 | E_refine | E_score2 |
| L<sub>(Native)</sub>(L<sub>co-</sub><br>crystalization | -5.8036 | 1.8487 | -<br>238.5506 | -129.0940 | -14.0081 | -5.1424 | -5.8036 |
| Complex1 | -4.5406 | 2.2353 | 44.0829 | -47.9389 | -10.2329 | -6.7920 | -4.5406 |
| Complex2 | -3.6741 | 1.4096 | 42.4540 | -48.6576 | -9.9953 | -6.3129 | -3.6741 |
| Complex3 | -4.9204 | 1.0998 | 99.2180 | -49.5140 | -12.0098 | -3.2896 | -4.9204 |
| Complex4 | -5.5403 | 2.3064 | 40.5023 | -34.6909 | -10.4948 | -12.0010 | -5.5403 |
| Complex5 | -4.7096 | 1.2774 | 48.5236 | -55.3976 | -10.0801 | -8.8089 | -4.7096 |
| Complex6 | -4.7270 | 2.2761 | 62.9089 | -38.9983 | -11.6355 | -11.1844 | -4.7270 |
| Complex7 | -4.9733 | 1.5392 | 42.9151 | -44.5609 | -10.3379 | -9.6226 | -4.9733 |
| Complex8 | -4.2295 | 2.5806 | 57.8758 | -38.4758 | -9.5209 | -12.3114 | -4.2295 |
| Complex9 | -5.9214 | 1.8308 | 86.4436 | -40.4958 | -12.3760 | -12.4170 | -5.9214 |
| Complex10 | -5.6293 | 1.0878 | 62.4814 | -56.2147 | -9.7901 | -13.5924 | -5.6293 |
| Complex11 | -5.4651 | 1.4103 | 73.9665 | -61.5321 | -11.6289 | -16.0503 | -5.7030 |
| Complex12 | -5.5836 | 1.7450 | 71.5921 | -53.9353 | -10.2497 | -14.9913 | -5.5836 |
| Complex13 | -5.4446 | 1.7018 | 65.1000 | -27.0714 | -9.7169 | -13.0686 | -5.4446 |
| Complex14 | -5.4800 | 2.5702 | 59.2881 | -41.4368 | -9.8105 | -13.6951 | -5.4800 |
| Complex15 | -4.4261 | 2.6518 | 44.6742 | -38.8736 | -10.4226 | -11.4980 | -4.4261 |
| Complex16 | -5.3617 | 1.9127 | 65.4036 | -67.6759 | -11.6111 | -8.9373 | -5.3677 |
| Complex17 | -6.1247 | 3.7014 | 93.0143 | -58.7966 | -10.5160 | -12.9499 | -6.1247 |
| Complex18 | -5.8086 | 2.0605 | 106.9323 | -66.7254 | -11.3439 | -6.0021 | -5.8086 |
| Complex19 | -5.3025 | 3.8129 | 95.5113 | -54.9084 | -10.9586 | -15.5251 | -5.3025 |
| Acarbose | -5.6234 | 4.7987 | 292.8790 | -121.9941 | -13.8717 | -23.2571 | -5.6234 |

### Binding interactions

The majority of active site residues found in our compounds have already been established in previous research, demonstrating the significance of these aminoacide residues in the active site of alpha-glucosidase **[39]**.

The figure **1** indicate clearly that L17, and Acarbose have a common amino acid residue MET 444 for binding in the active site of 3L4Y. In the same figure the ligands L4, L11, and Acarbose have the same amino acid residue ASP 542 in the pocket of alpha-glucosidase. ASP 542 was matched with the potential binding sites and favorable interacting residues of 3L4Y **[39]**.

**Fig 1.**
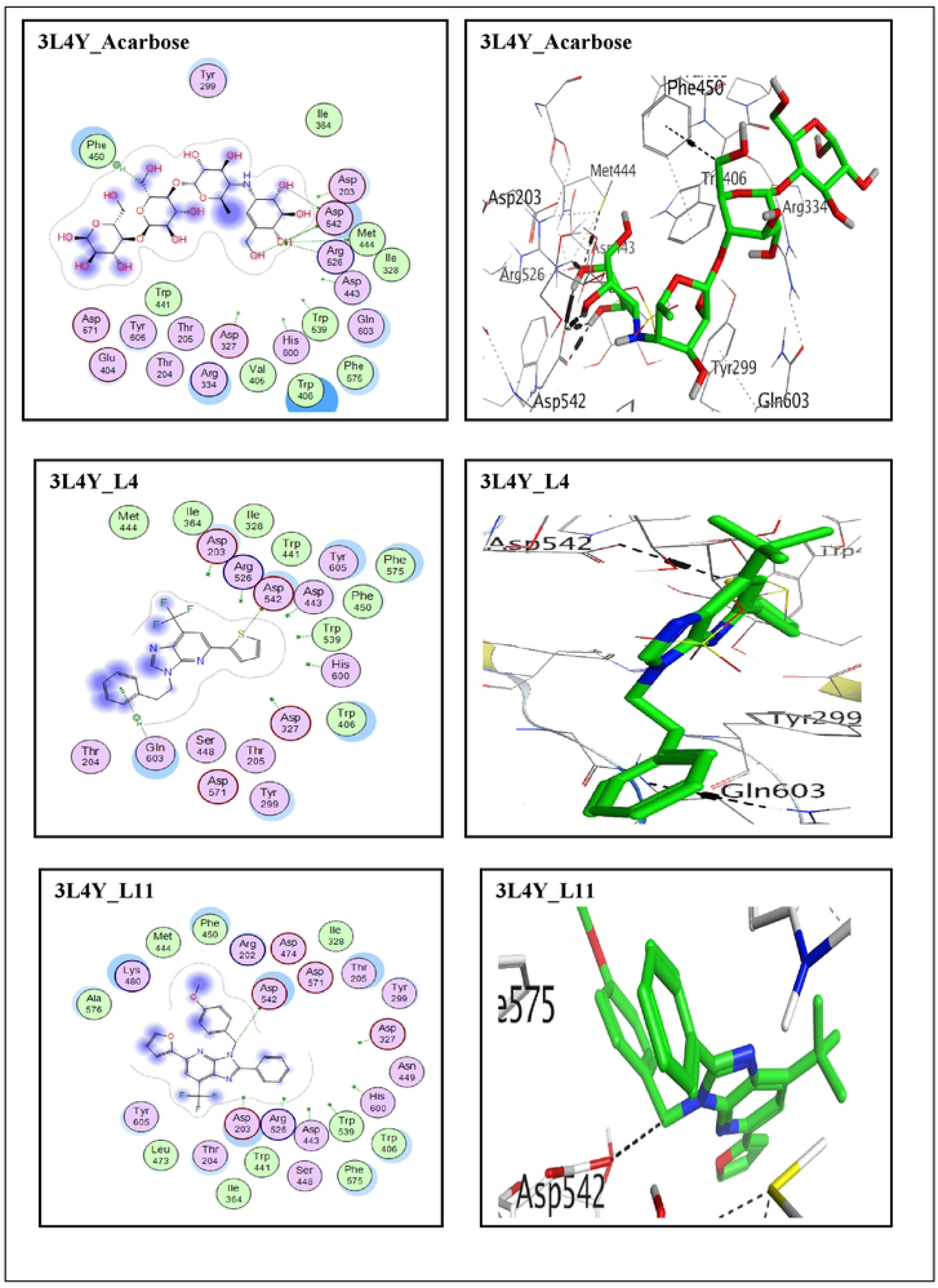
2D and 3D visualization of the best poses of **acarbose**, **L4**, **L11** and **L17** with site active of 3L4Y

### Evaluation of properties druglikeness

The Drug-likeness set is composed of ligands that satisfy “Lipinski’s Rule of Five” **[43]**, which are four chemical properties of compounds that are suitable for drugs **[44]**, molecular weight ≤ 500g/mol, MLogP < 5, number of hydrogen bond donor < 5, and hydrogen bond acceptors < 10. The results presented in table 3 indicated that all ligands complied with Lipinski’s rule with exception of Acarbose. In addition estimation of properties such as the partition coefficient (−2 ≤ LogP ≤ + 6) and polar surface area (TPSA: less than 140Å) **[45]** can be used as a preliminary filter to remove compounds that are likely to show permeability and solubility problems and, consequently, poor pharmacokinetic profiles **[43]**. We note that the good result of TPSA is observed for the ligands but not for Acarbose (table 4).

**Table 3.** Results of bonds between atom of ligand and residue of active site into 3L4Y of molecular docking.

| Name of ligands | Atom of the ligand | Involve receptor residues | Types of interaction bonds | Distance (Å) |
| --- | --- | --- | --- | --- |
| **L4** | S | ASP 542 | H-donor | 3.55 |
|  | 6-ring | GLN 603 | Pi-H | 4.46 |
| **L6** | / | / | / | / |
| **L11** | C | ASP 542 | H-donor | 3.61 |
| **L12** | / | / | / | / |
| **L13** | / | / | / | / |
| **L17** | C | MET 444 | H-donor | 3.88 |
|  | O | THR 205 | Pi-H | 3.10 |
| Acarbose | O | ASP 542 | H-donor | 3.06 |
|  | O | MET 444 | H-donor | 3.69 |
|  | O | ASP 542 | H-donor | 2.77 |
|  | O | ASP 542 | H-donor | 3.11 |
|  | O | ASP 542 | H-donor | 3.16 |
|  | O | ARG 526 | H-acceptor | 2.89 |
|  | C | PHE 450 | Pi-H | 4.37 |

**Table 4.**
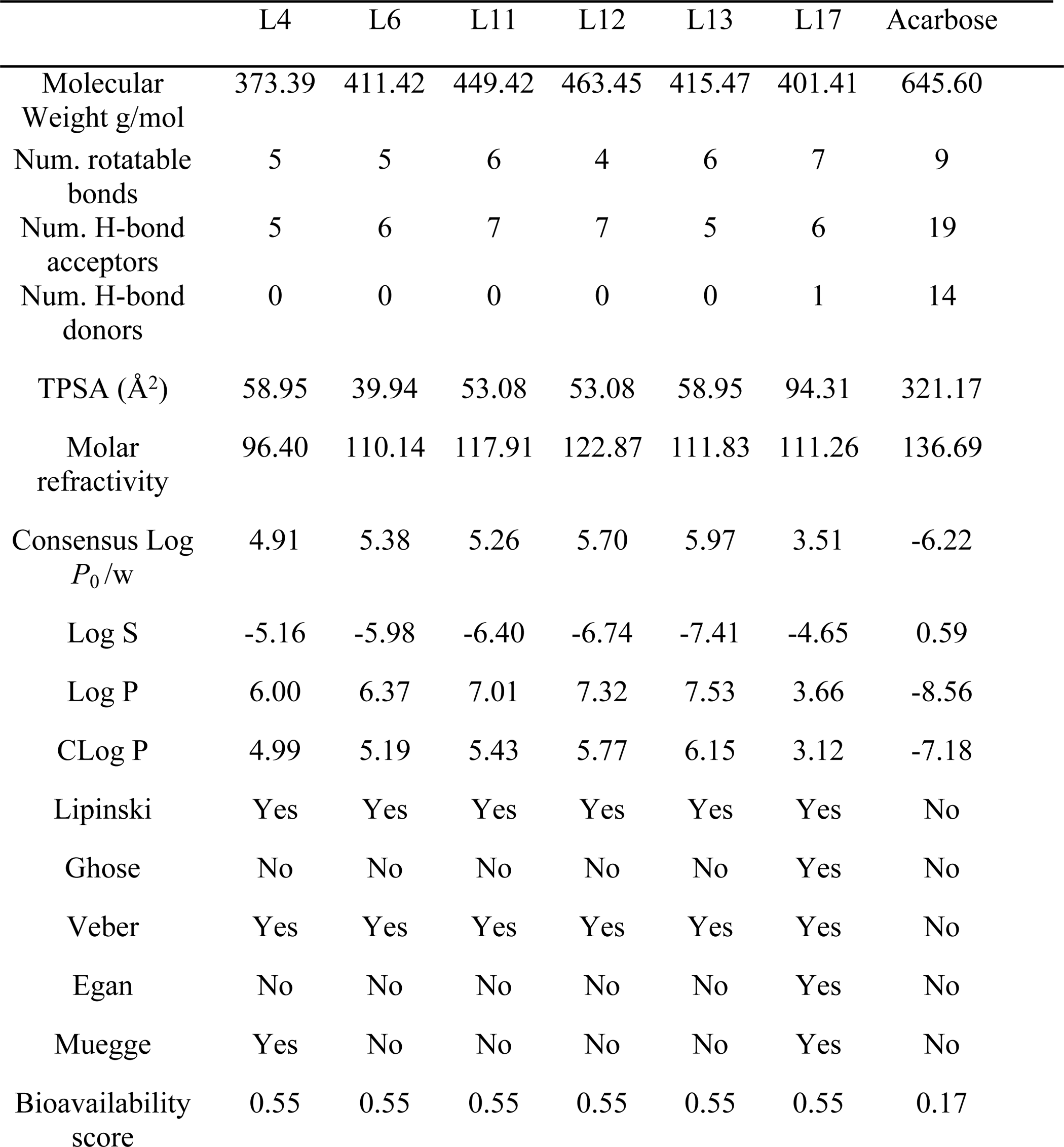
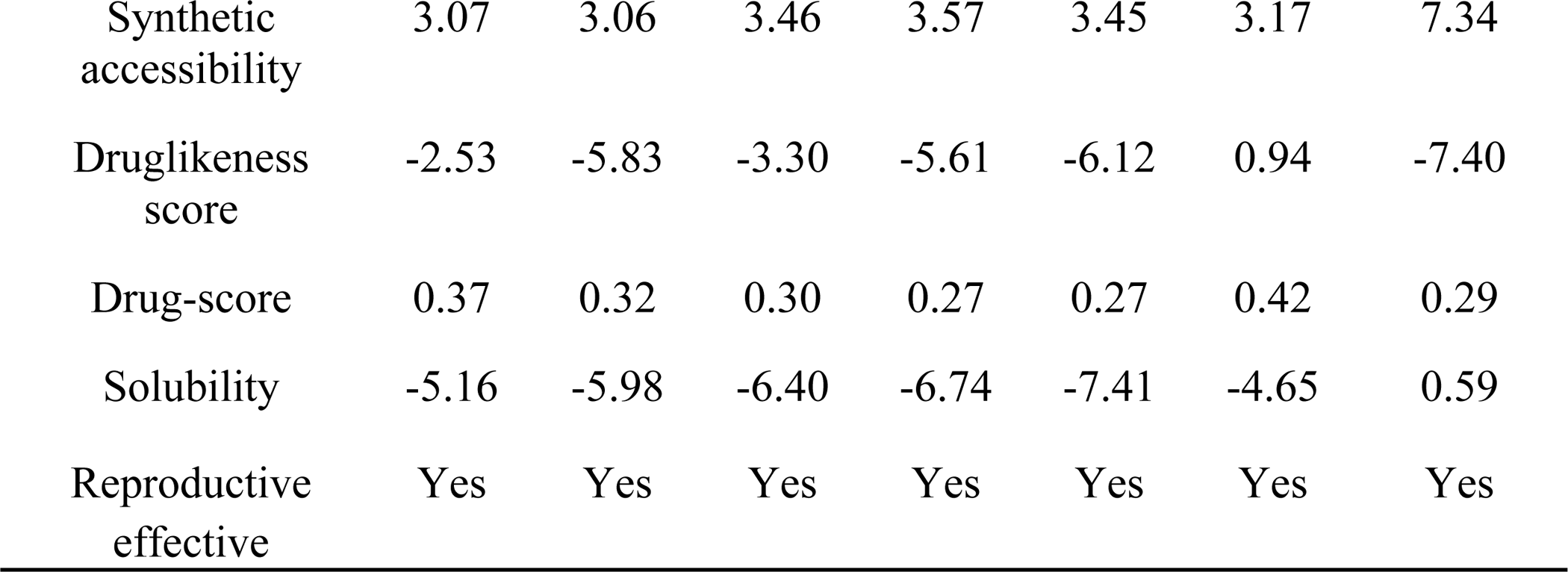
Druglikeness properties of the main ligands.

### ADMET properties

ADMET filters are integrated in the early stages of drug discovery and development. Absorption depends on the triad “potency-permeability-solubility” **[46]**.

The inclusion of in silico in vitro absorption, distribution, metabolism, excretion, and toxicity (ADMET) filters in the early stages of drug discovery **[47,48]** and the emergent interest in low affinity ligands within certain therapeutic categories **[49]**.

The results of the ADMET test are showed in the table 5. For the pKCSM predictive model high CaCo_2_ permeabitily would translate in predicted values > 0.90. Compounds L4 and L17 have a high CaCo_2_ permeabitily. Moreover the same ligands have higher HIA than 30%; suggesting that these ligands can be administered orally and will be absorbed through the human intestine. We can see also that these ligands L4, L11 and L17 showed average skin permeability because their Log Kp is slightly higher to −2.5. Concerning the distribution, the compound who present Log BB > 0.3 can cross the blood brain barrier, it is not in our case. For the compound who has Log PS > −2, we can said that L4 and L11 is able to penetrate the CNS.

**Table 5.** ADMET test results by PkCSM.

| Property | Model name | Predict value |  |  |  |
| --- | --- | --- | --- | --- | --- |
|  |  | Acarbose | L17 | L4 | L11 |
| Absorption | Water solubility | -1.482 | -2.981 | -3.115 | -2.893 |
| Absorption | CaCo<sub>2</sub> permeability | -0.481 | 1.13 | 0.422 | -0.581 |
| Absorption | Intestinal absorption human (HIA) | 4.172 | 100 | 89.212 | 86.992 |
| Absorption | Skin permeability | -2.735 | -2.735 | -2.738 | -2.735 |
| Absorption | P-glycoprotein substrate | Yes | Yes | No | Yes |
| Absorption | P-glycoprotein I inhibitor | No | Yes | Yes | Yes |
| Absorption | P-glycoprotein II inhibitor | No | Yes | Yes | Yes |
| Distibution | BBB permeability | -1.717 | -0.648 | 0.25 | 0.47 |
| Distibution | CNS permeability | -6.438 | -3.377 | -1.303 | -1.68 |
| Metabolism | CYP2D6 substrate | No | No | No | No |
| Metabolism | CYP3A4 substrate | No | No | Yes | No |
| Metabolism | CYP1A2 inhibitor | No | Yes | Yes | Yes |
| Metabolism | CYP2C19 inhibitor | No | Yes | Yes | Yes |
| Metabolism | CYP2C9 inhibitor | Yes | No | No | Yes |
| Metabolism | CYP2D6 inhibitor | No | No | Yes | No |
| Metabolism | CYP3A4 inhibitor | No | Yes | No | Yes |
| Excretion | Total clearance | 0.435 | 0.906 | 0.849 | 0.396 |
| Toxicity | AMES Toxicity | No | No | Yes | No |
| Toxicity | Hepatotoxicity | No | No | No | No |
| Toxicity | Skin sensitisation | No | No | No | No |

### Molecular Dynamics (MD) simulation

Molecular dynamics simulation offers a robust toolbox that allows us to delve into the intricate details of structure and dynamics **[32]**. This approach proves particularly valuable in introducing protein flexibility post a docking protocol, refining the structures of protein–drug complexes **[43]**.

The outcomes of the molecular dynamics (MD) simulation for the L17, L11, and L4 complexes align with the findings from molecular docking tests. Throughout the simulation trajectory, various metrics, including the radius of gyration, solvent accessible surface area (SASA), root mean square fluctuations (RMSF), total number of hydrogen bonds, and root mean square deviation (RMSD), were examined.

Initially, the RMSD plot indicated deviations in every complex during a certain time period. Subsequently, it was observed that the complex containing L11 remained stable throughout the entire simulation, extending up to 100 ns. The data revealed that the RMSD of various protein-matched ligands varied between 0.12 and 0.21 nm with occasional peaks reaching 0.23 nm. Notably, the RMSD of the protein-matched ligand L11 exhibited a consistent, steady graph. Between 40 and 60 ns, the ligand-fitted protein’s RMSD fluctuated between 0.15 and 0.23 nm, stabilizing after 60 ns. These observations strongly suggest the formation of a stable complex between L11 and alpha-glucosidase, facilitated by favorable interactions with crucial amino acid residues (Fig 2).

**Fig 2.**
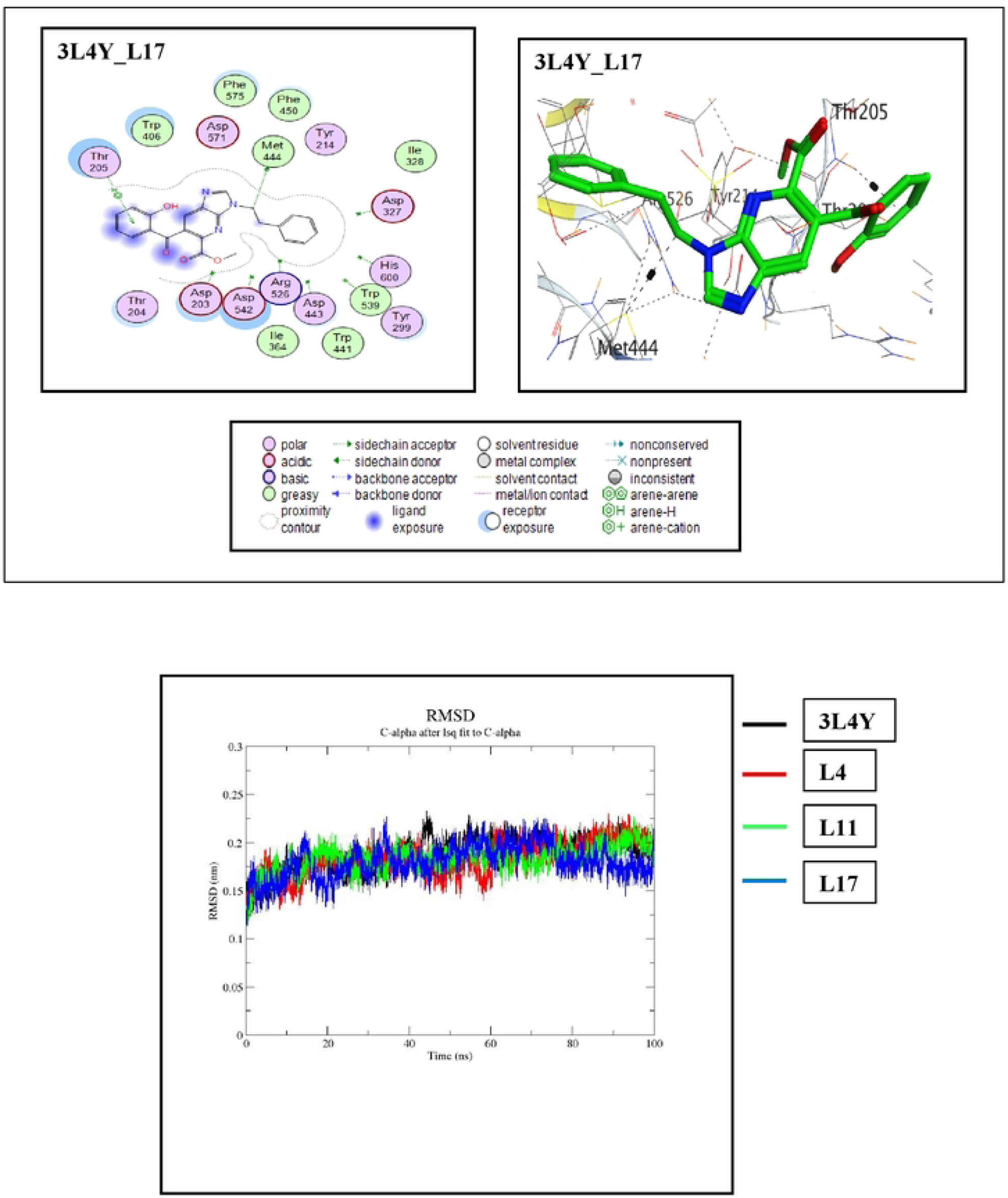
RMSD Plot for 3L4Y complex with L4(red), L11(green), L17(blue) and 3L4Y alone (black)

RMSF trajectories provide vital insights into the stability of a complex. High fluctuations indicate increased flexibility and unstable bonds, while low values signify less distortion and well-structured areas within the complex. Fig. 3 illustrates that the patterns exhibited by each system were nearly identical. The overall RMSF value for each of the three complexes, as observed in Fig. 3 was approximately 0.3 nm.

**Fig 3.**
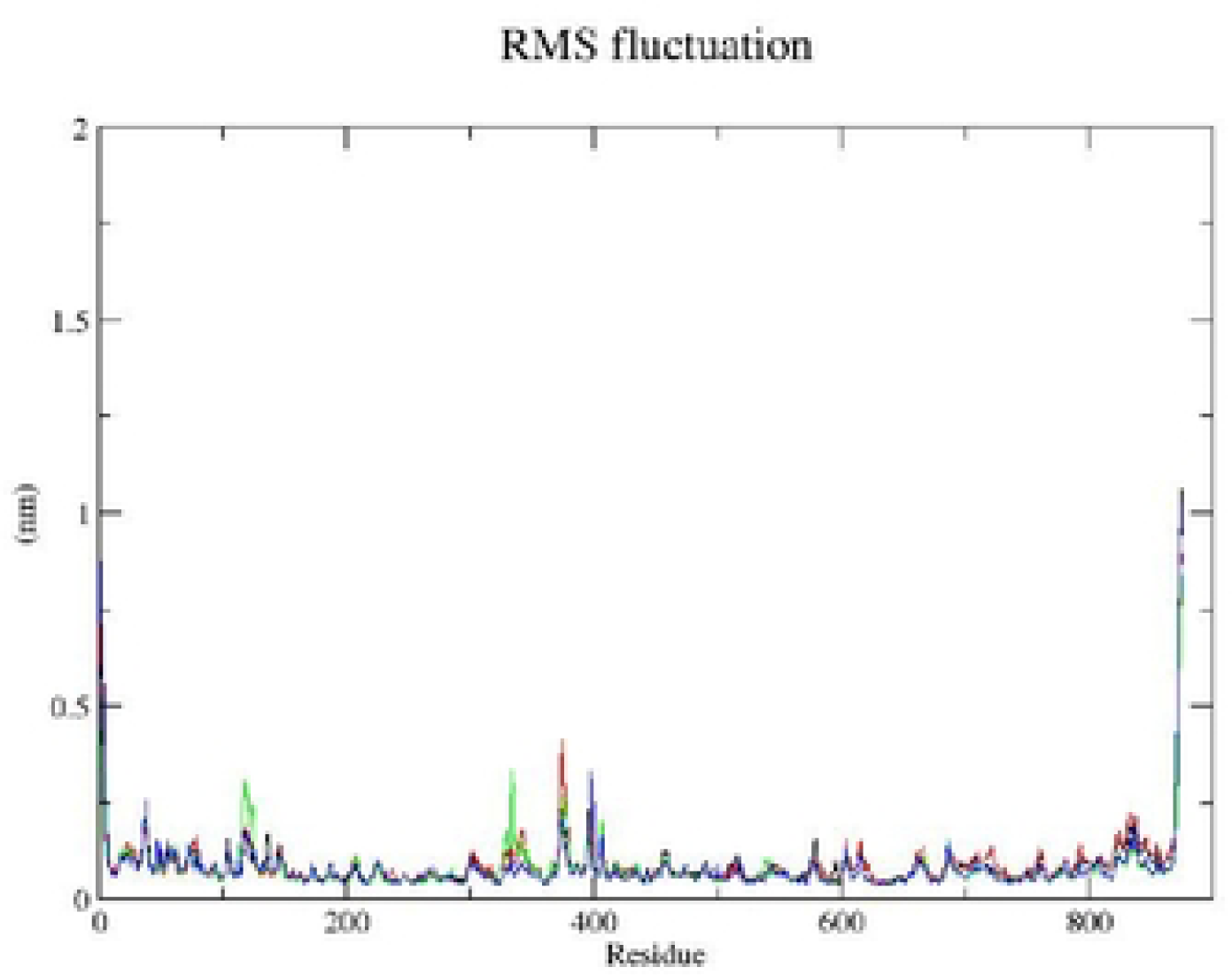
RMSF plot for 3L4Y complex with L4(red), L11(green), L17(blue) and 3L4Y alone (black)

Plotting Rg against time was performed to assess the system’s compactness. Lower Rg values indicate good structural stability (more folded), while higher Rg values suggest less compactness with conformational entropy. The simulation Rg values for the three compounds range between 0.289 and 0.292 nm, as shown in Fig 4.

**Fig 4.**
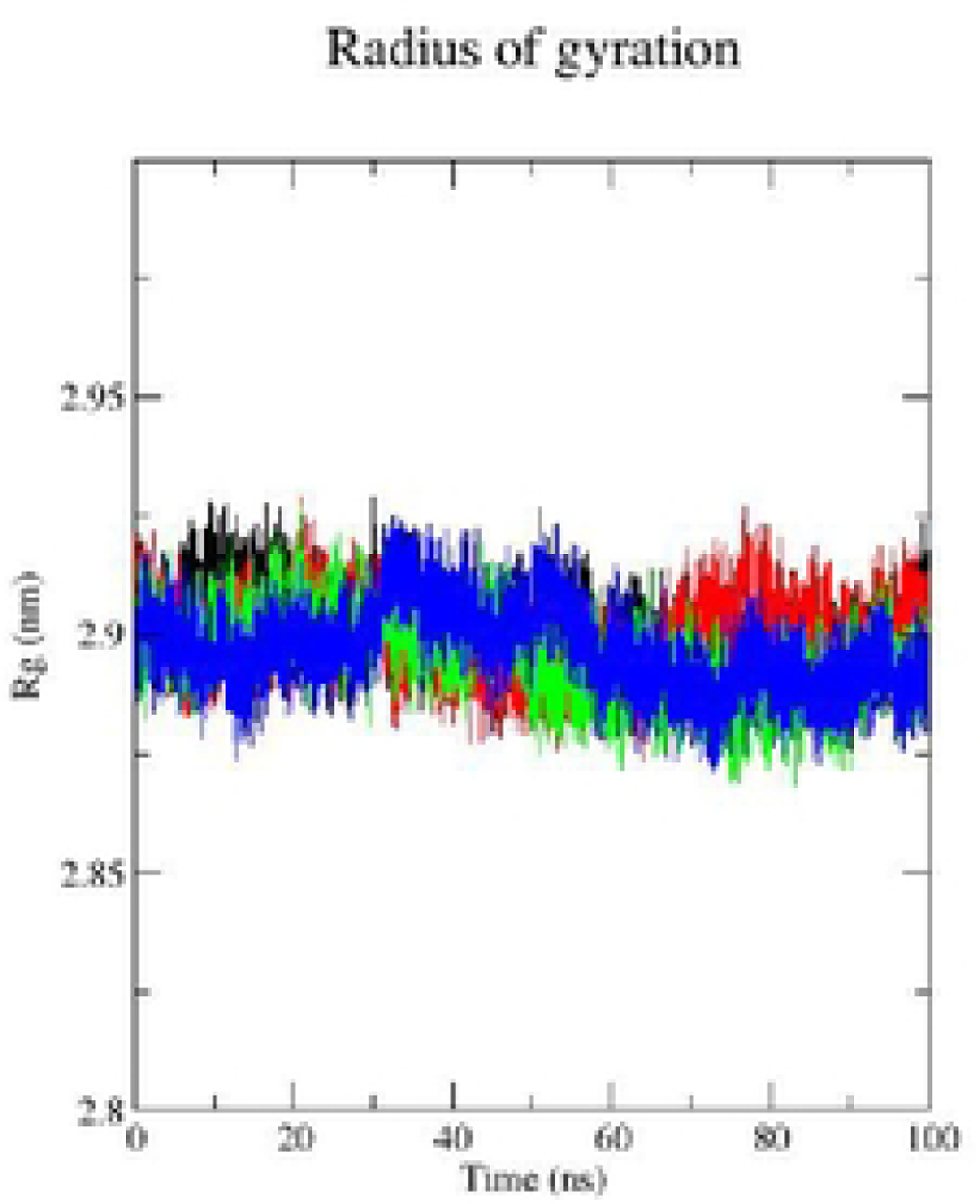
RMSF plot for 3L4Y complex with L4(red), L11(green), L17(blue) and 3L4Y alone (black)

Consistent Rg values indicate stability within the protein complex, suggesting that the binding of these three molecules does not induce significant structural alterations. The compact architecture and size of the three complexes are substantiated by their respective Rg ranges, as illustrated in Fig 4.

The degree to which ligands and proteins bind is determined in large part by hydrogen bonding. Throughout the simulation, L11 (green) id (black) has a consistent hydrogen bond range of 0 to 2, whereas L4 displayed bonding variations. There are more hydrogen(>4) bonds between 30 and 65 ns, which could indicate that the binding site’s conformation changed throughout simulation (Fig 5). Overall, the observations indicated that alpha glucosidase is stable during simulation when complexed with the three chemicals.

**Fig 5.**
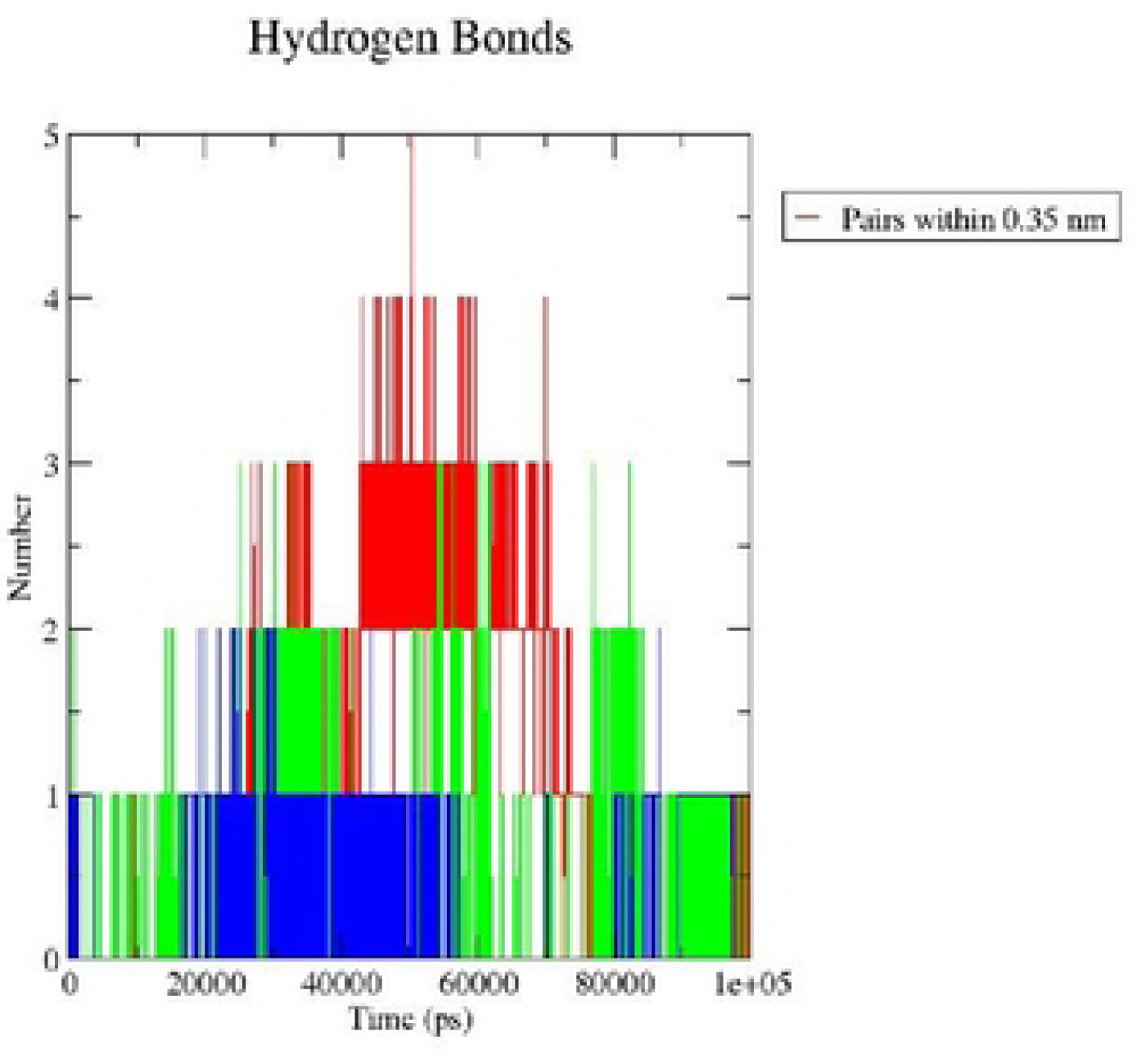
Hydrogen bonds plot for 3L4Y complex with L4(red), L11(green), L17(blue) and 3L4Y alone (black)

On the other hand the solvent-accessible surface area of proteins (SASA) is a measure that reveals information on the relative exposure or burying of these molecules inside the protein structure.

The SASA plots of the alpha-glucosidase complexes in the figure exhibit variations during the simulation. Each complex demonstrates a consistent downward trend in SASA values over time. Notably, the enzyme-ligand complexes display higher SASA values, suggesting increased exposure to solvent molecules compared to unbound alpha-glucosidase.

Interestingly, among the complexes, ligand L11 stands out with the lowest SASA value, indicating a more compact or buried shape with minimal solvent exposure. This suggests the possibility of tighter binding or stronger interactions between L11 and alpha-glucosidase, resulting in a more restricted and less exposed complex of enzyme and ligand (Fig 6).

**Fig 6.**
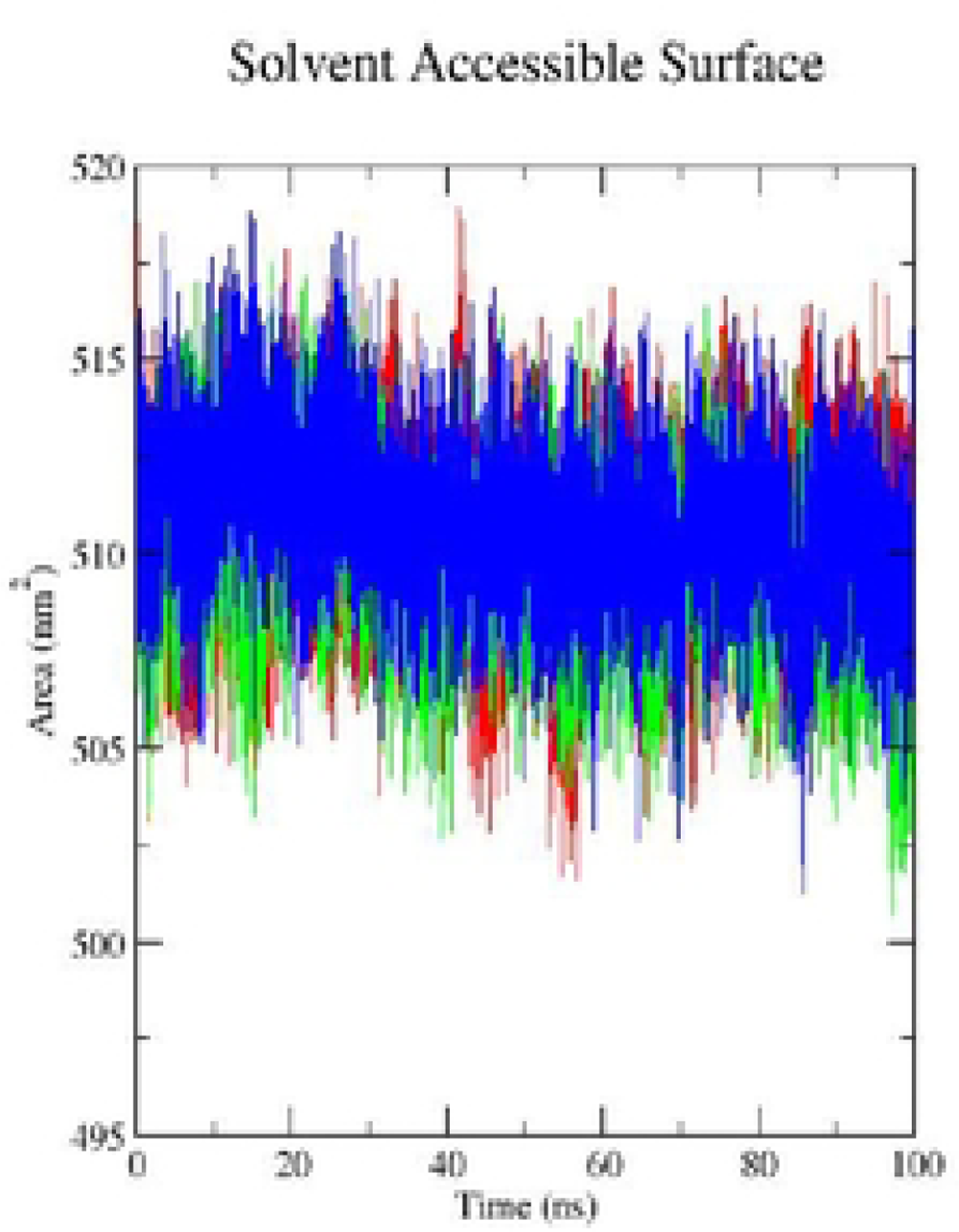
Solvent accessible surface area for the 3L4Y and the studied complexes over the simulation time

## Conclusion

A series of 1-deazapurines derivatives have already been synthesized and tested for their ability to inhibit alpha-glucosidase. Six of them (Table 1) were shown to inhibit the alpha-glucosidase enzyme with a higher percentage of inhibition and lower IC50 values. In this study, the understanding profile of binding of these bioactive chemicals to alpha-glucosidase is predicted, and it is contrasted with that of Acarbose, a known active inhibitor by molecular docking. The ligands L17, L11, and L4 have the best binding energy among those compounds with establishing interaction into the active site. In addition MD simulation is used to optimize the molecular docking results under dynamic conditions this shows that the best ligands are stable. Furthermore, ADME-T predictions highlight that L17 is the only compound among the 19 tested that meets all drug-likeness requirements without violating Lipinski, Ghose, Veber, Mueggy, or Egan rules. On the other hand, L4 and L11 crosses the central nervous system. For the role of an alpha-glucosidase inhibitor in the anti-T2DM drug candidate, L11 and L4 emerge as promising multi-target drug candidates as indicated by dynamic simulations.

## Supporting information

**S1 Fig.** 2D and 3D visualization of the best poses of **acarbose**, **L4**, **L11** and **L17** with site active of 3L4Y

**S2 Fig.** RMSD Plot for 3L4Y complex with L4(red), L11(green), L17(blue) and 3L4Y alone (black)

**S3 Fig.** RMSF plot for 3L4Y complex with L4(red), L11(green), L17(blue) and 3L4Y alone (black)

**S4 Fig.** RMSF plot for 3L4Y complex with L4(red), L11(green), L17(blue) and 3L4Y alone (black)

**S5 Fig.** Hydrogen bonds plot for 3L4Y complex with L4(red), L11(green), L17(blue) and 3L4Y alone (black)

**S6 Fig.** Solvent accessible surface area for the 3L4Y and the studied complexes over the simulation time

**S1 Table.** Structure of ligands and their respective percentage of inhibition and IC_50_

**S2 Table**. Energy score of 19 ligands of 1-deazapurines

**S3 Table.** Results of bonds between atom of ligand and residue of active site into 3L4Y of molecular docking

**S4 Table.** Druglikeness properties of the main ligands

**S5 Table.** ADMET test results by PkCSM

## Acknowledgments

We are grateful to the group of computational and biological activity team, LASNABIO Laboratory of University of Tlemcen - Algeria.

## Funding

This research received no external funding. No funding was received to assist with the preparation of this manuscript.

## Declaration of Competing Interest

The authors declare that they have no known competing financial interests or personal relationships that could have appeared to influence the work reported in this paper

## Notes

### Competing Interest Statement

The authors have declared no competing interest.

